# The semantic segmentation approach for normal and pathologic tympanic membrane using deep learning

**DOI:** 10.1101/515007

**Authors:** Jungirl Seok, Jae-Jin Song, Ja-Won Koo, Hee Chan Kim, Byung Yoon Choi

## Abstract

**Objectives:** The purpose of this study was to create a deep learning model for the detection and segmentation of major structures of the tympanic membrane.

**Methods:** Total 920 tympanic endoscopic images had been stored were obtained, retrospectively. We constructed a detection and segmentation model using Mask R-CNN with ResNet-50 backbone targeting three clinically meaningful structures: (1) tympanic membrane (TM); (2) malleus with side of tympanic membrane; and (3) suspected perforation area. The images were randomly divided into three sets – taining set, validation set, and test set – at a ratio of 0.6:0.2:0.2, resulting in 548, 187, and 185 images, respectively. After assignment, 548 tympanic membrane images were augmented 50 times each, reaching 27,400 images.

**Results:** At the most optimized point of the model, it achieved a mean average precision of 92.9% on test set. When an intersection over Union (IoU) score of greater than 0.5 was used as the reference point, the tympanic membrane was 100% detectable, the accuracy of side of the tympanic membrane based on the malleus segmentation was 88.6% and detection accuracy of suspicious perforation was 91.4%.

**Conclusions:** Anatomical segmentation may allow the inclusion of an explanation provided by deep learning as part of the results. This method is applicable not only to tympanic endoscope, but also to sinus endoscope, laryngoscope, and stroboscope. Finally, it will be the starting point for the development of automated medical records descriptor of endoscope images.

## Introduction

Otoscopic examination is one of the most basic screening tools for checking the condition of the ear canal and tympanic membrane. Acute otitis media (AOM), chronic otitis media (COM), otitis media with effusion (OME), cholesteatoma, and perforation of the tympanic membrane can easily be diagnosed by simple tympanic membrane examination and clinical presentation.

Otolaryngologists, pediatricians, and general practitioners perform otoscopic examinations on a daily basis as a part of routine care [1]. As with other medical tests, there is an observer-to-observer variation in the accuracy of the diagnosis, depending on the technical skill and experiences of the physician as well as observer’s subjective bias. To improve the accuracy, so-called computer-aided detection or diagnosis (CAD) may be required.

CAD is a computerized procedure that provides an objective interpretation of the medical image [2]. With the recently introduced deep learning technique, promising results have been obtained in the CAD field [2]. However, these results were obtained mostly based on radiologic tests, such as CT and MRI, or ophthalmologic tests. Studies using endoscopic images are rarely conducted.

To date, there have been a few studies regarding otoscopic examination using deep learning or machine learning techniques; however, they only showed a simple binary classification result, either normal or abnormal findings [3–5]. A simple classification may not be sufficient to be applicable in clinical practice. For example, a simple, ‘abnormal’ classification may be too broad, not providing the ‘why’ and ‘where’ of the problem. To overcome these shortcomings, it is necessary to create a deep learning model that provides better interpretable and informative results by dividing and segmenting the whole image into sets or parts based on detailed structure or findings.

The purpose of this study was to create such a deep learning model for the detection and segmentation of major structures of the tympanic membrane and to verify the applicability of the model.

## Materials and Methods

### Data Set

The study protocol was reviewed and approved by the Institutional Review Board of Seoul National University Bundang Hospital, and a waiver of informed consent was granted because the study utilized images that had been stored; the method of data collection was the same as that of retrospective design study (IRB No. B-1808/489-104).

For model development, tympanic endoscopy images, which had been previously taken from 2004 through 2017, were obtained from Seoul National University Bundang Hospital. Personal information was excluded when extracting images from the PACS system.

To create a universal algorithm applicable to any clinical setting, the images were extracted randomly, except the ones that were too blurry to identify. All images were taken from various locations (outpatient clinics or wards) with various specifications of charge-coupled device (CCD) and endoscope. There was a limitation in terms of acquiring the exact setting on the endoscopy at the time of the tympanic membrane photograph. To avoid redundant training of similar tympanic membrane images, only one image was obtained from each ear of each patient. A total of 920 images were obtained; the images were randomly divided into three sets – training set, validation set, and test set – at a ratio of 0.6:0.2:0.2, resulting in 548, 187, and 185 images, respectively.

### Labeling

Labeling of all images was done by one skilled otolaryngologist using the *labelme* application (https://github.com/wkentaro/labelme). Another experienced otolaryngologist reviewed and verified the labeling (**Fig 1**). The label was then used as a mask of the structure.

**Fig 1.**
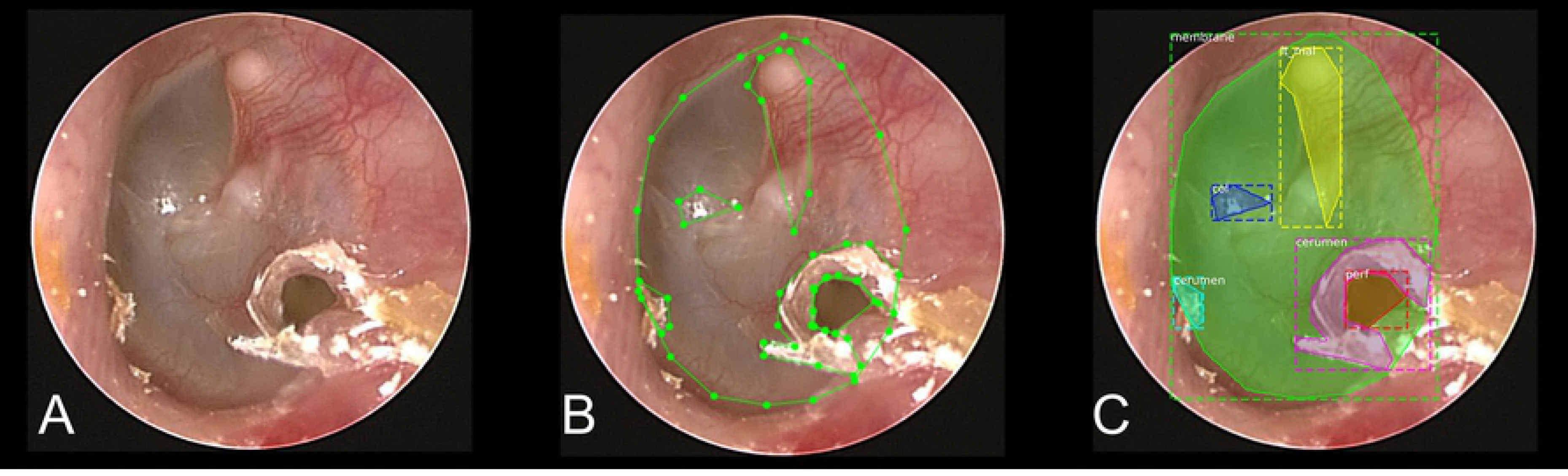
Original tympanic endoscopy image (A), labeled image (B), and visualized labels of image (C).

The labeled structures were the tympanic membrane (TM), malleus, suspected areas of perforation, cone of light, cerumen, and tympanosclerotic plaque (TSP). In the case of the malleus, it was labeled in association with the side of the tympanic membrane due to the side classification (left vs right) of the tympanic membrane, which can be achieved by the shape of the malleus neck or the location of the head of the malleus. If the malleus was not visible, the direction of the tympanic membrane was classified as ‘undefined’. And perforation was not completely distinguishable from severe retraction, because this study was performed but on a still image, not a moving picture. Therefore, such findings were all defined as ‘suspicious perforation’.

In the course of the study, labels of cone of light, cerumen, and TSP were finally excluded from the final model due to the lack of clinical significance (**S1 Fig**).

### Development of the algorithm

Mask R-CNN [6] implementation by Matterport, Inc (Mountain View, CA, USA) (https://github.com/matterport/Mask_RCNN) with ResNet-50 [7] backbone was used. The model was modified for the purpose of our study. The deep learning model is able to learn by itself through the tympanic membrane images that contain the pre-labeled regions, and then it is able to find the region corresponding to each anatomical structure, with the input of the tympanic membrane images.

Because training the model requires a large number of images, 548 tympanic membrane images were augmented 50 times each, reaching 27,400 images. This augmentation was obtained by the following methods: (1) horizontal flip; (2) noise addition; (3) Gaussian blur (σ: 0.0-2.0); (4) rotation using center coordinates of a tympanum; and (5) zoom in and out of images. (**S2 Fig**) These methods were implemented using the *imgaug* library (https://github.com/aleju/imgaug). Horizontal flip was performed with a probability of 50%, and other augmentation methods were executed randomly.

Endoscope images are usually all black, except the middle-rounded area. These black areas of the image are unnecessary. After applying the blur effect, only the rounded, meaningful regions were extracted using both the Otsu threshold method and region detection based on the adaptive threshold. Following this, the long axis was adjusted to 512 pixels, and finally a 512×512 pixel image was obtained by adding the padding. (**S3 Fig**) The masks drawn along the boundary of the structures were dilated to 20 pixels, which could allow the boundary line itself to be a feature of the structure.

The weights of the model were initialized using a pre-trained model, which was the same network as those trained with the MS-COCO data set [8].

The trained model generates an array of class indexes of the structure; for example, the tympanic membrane is index 1, the right malleus is index 2, and the left malleus is index 3. Then the model creates a mask that represents the area of the structure. In other words, each value of the array is the class index, and this value has a corresponding mask that represents the position and size of the region.

### Evaluation of the Algorithm

As the training continued, the loss value was obtained by using both the training and validation data sets. If the training loss value continues to decrease and the validation loss value starts to increase, it can be considered as overfitting from that point onward. At this point, the actual test with test set images was carried out after selecting the model. The results were analyzed by dividing the images into individual structures, and we evaluated the detection rate and segmentation accuracy using the intersection over union (IoU) score. In general, an IoU score of over 0.5 is normally considered as a good prediction in semantic segmentation research [9].

## Results

At the most optimized point of the model, it achieved a mean average precision (mAP) of 92.9% on test set. The actual output result using the learned model is shown in **Fig 2**. Accurate recognition was achieved. In the case of large perforation, all the possibilities, including erroneous recognition due to neighboring structures visualized through the perforation, were all provided. (**Fig 3**)

**Fig 2.**
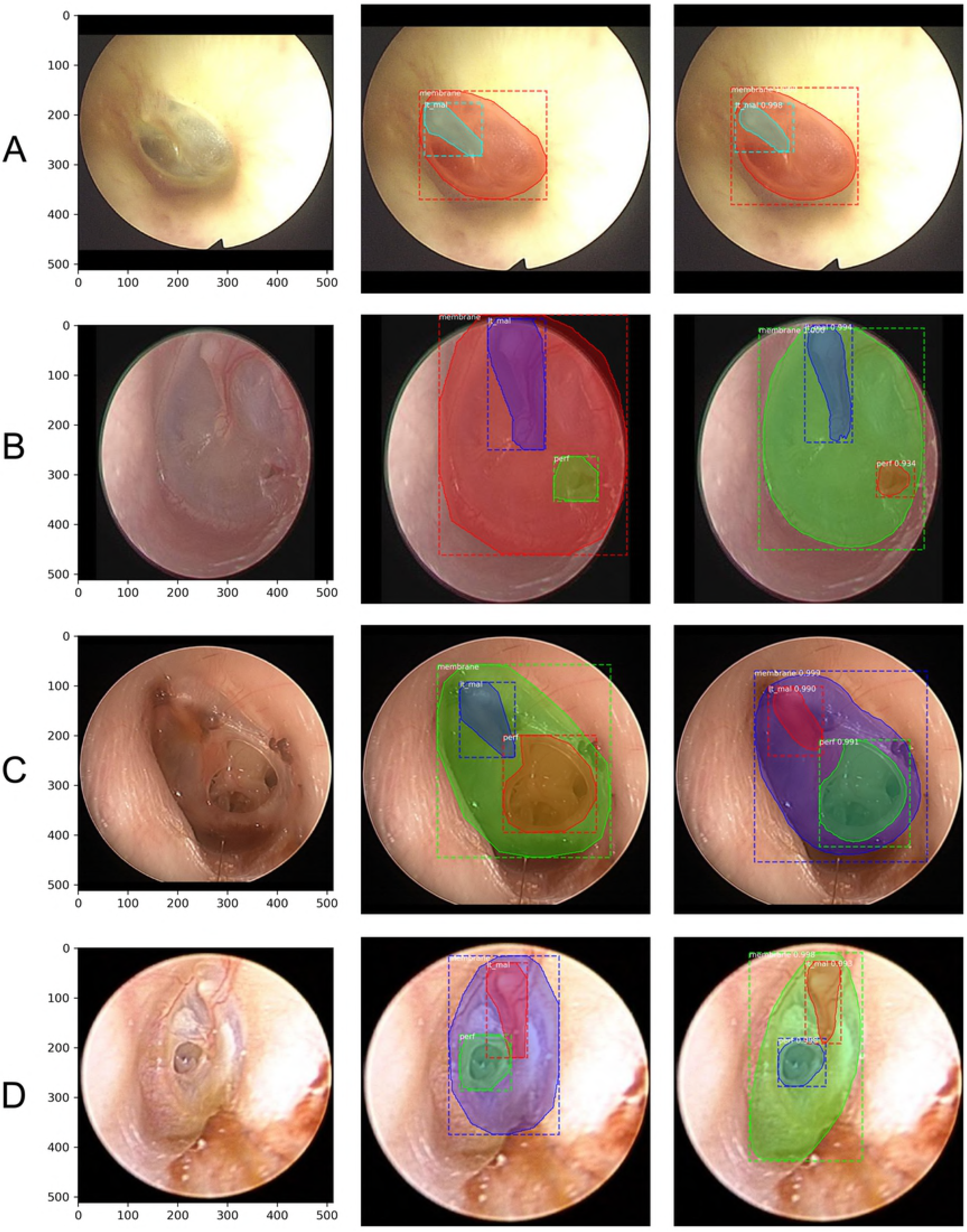
Original images (left), ground truth (center), and outputs (right). The tympanic membrane, malleus with orientation, and perforation area are successfully detected and segmented. The label, prediction scores calculated under the model and segmented regions are provided in the outputs.

**Fig 3.**
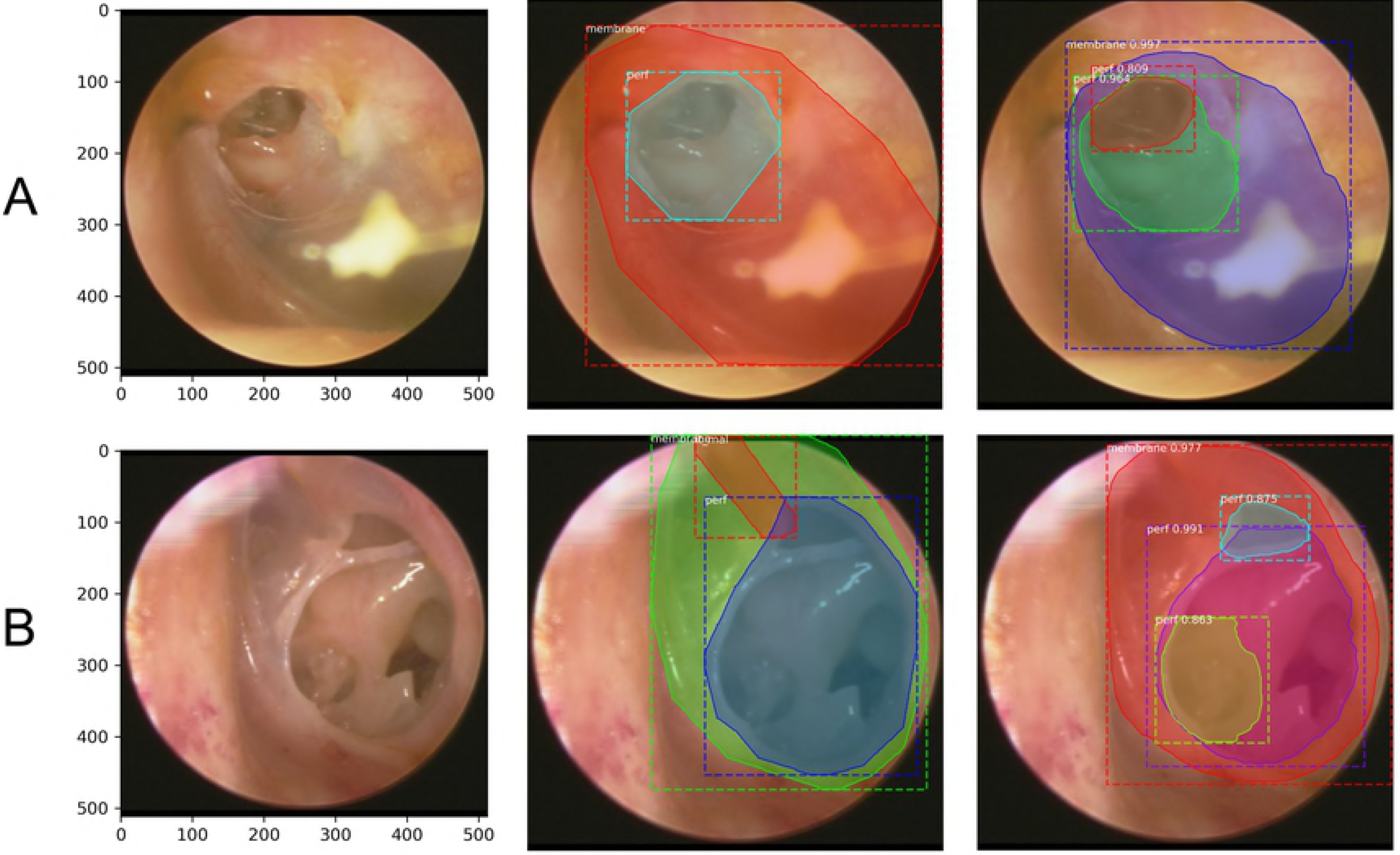
Results obtained from TM images with large perforation as shown by original images (left), ground truth (center), and outputs (right). (A,B) Inside each large perforation, other structures or a part of the true structures are erroneously recognized also as small perforations due to the neighboring structures. The malleus is not detected in the both output images, but in the Fig 3A, malleus is hard to define in the ground truth image (true negative), but in Fig 3B, malleus is visible in the ground truth (false negative).

### Accuracy of TM segmentation

In the images with low IoU scores, for example, 0.5 indicates that only about half of the area is detected. In images with an IoU score of greater than 0.5, the tympanic membrane was 100% detectable in deep learning and the detection rate still remained over 95% when an IoU score is over 0.7. (**Table 1**).

**Table 1.** The segmentation rate of the tympanic membrane, which meets the criteria according to IoU values, respectively

|  | IoU |  |  |  |
| --- | --- | --- | --- | --- |
|  | > 0.5 | > 0.6 | > 0.7 | > 0.8 |
| Successful segmentation rate (n=185) | 100% | 98.4% | 95.1% | 80.0% |

Specifically, in the actual image where it is difficult to delineate the exact boundary of the tympanic membrane, most of the important tympanic membrane area, including the perforation, were detected even when the IoU score was as low as 0.516 (**Fig 4A**). Our results showed that segmentation can be achieved with meaningful level of accuracy using deep learning, even with an IoU score of just over 0.6 (**Fig 4B, 4C**).

**Fig 4.**
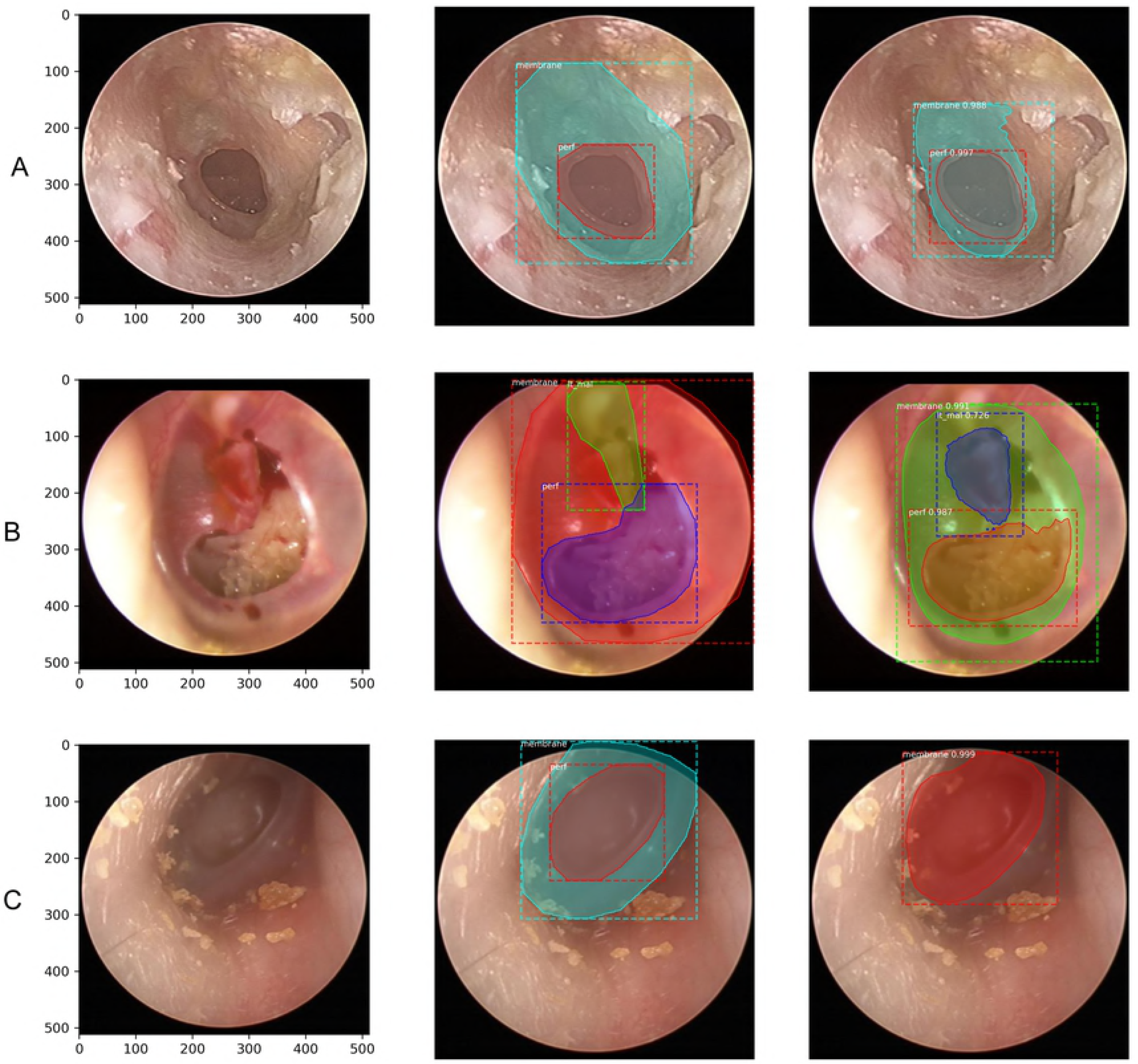
Exemplary images with low IoU values. (A) 0.516, (B) 0.651, and (C) 0.665. Original images (left), ground truth (center) and outputs (right). Fig 4A shows the tympanic membrane with difficulty in accurately defining the boundary even with expert’s eye and Fig 4B, C seems to contain enough normal tympanic membrane area even though low IoUs are obtained.

### Accuracy of malleus segmentation

The results of evaluating the side of the tympanic membrane (right vs left) based on the malleus segmentation are shown in **Table 2** and **Table 3**. If IoU was not considered, the accuracy was 95.1%, and with the consideration of an IoU score of greater than 0.5, the accuracy was 88.6%. Among the cases with IoU over 0.5, when the side was decided by the model, the agreement rate was 99.2%. (Positive predictive value).

**Table 2.**
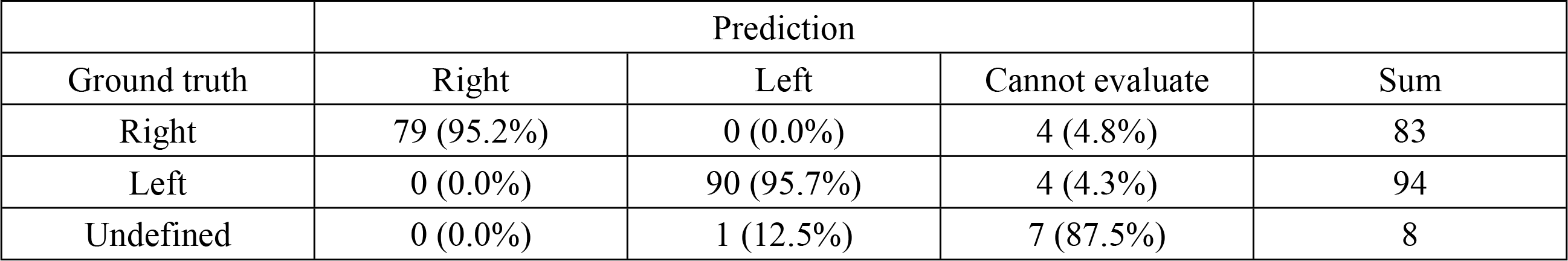
Results of malleus detection and side prediction using the model

**Table 3.**
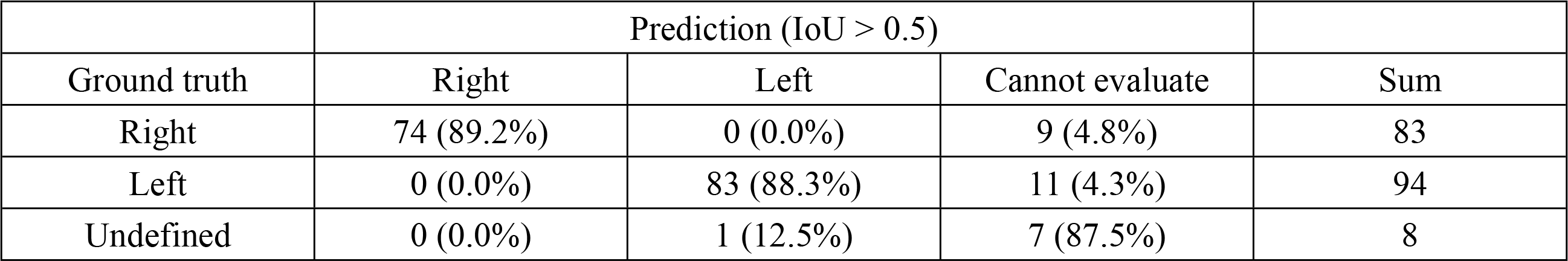
Results of malleus detection and side prediction using model when IoU > 0.5 was defined as valid

|  | Prediction (IoU > 0.5) |  |  |  |
| --- | --- | --- | --- | --- |
| Ground truth | Right | Left | Cannot evaluate | Sum |
| Right | 74 (89.2%) | 0 (0.0%) | 9 (4.8%) | 83 |
| Left | 0 (0.0%) | 83 (88.3%) | 11 (4.3%) | 94 |
| Undefined | 0 (0.0%) | 1 (12.5%) | 7 (87.5%) | 8 |

### Accuracy of perforation segmentation

Detection accuracy of suspicious perforation without considering the IoU score and at IoU > 0.5 was 93.0% and 91.4%, respectively (**Table 4, Table 5**). The results of perforation lesion detection can be seen in **Fig 2, Fig 3,** and **Fig 4.**

**Table 4.** Detection results of suspected area of perforation.

|  | Prediction |  |  |
| --- | --- | --- | --- |
| Ground truth | Suspicious perforation | Intact | Sum |
| Suspicious perforation | 51 (86.4%) | 8 (13.6%) | 59 |
| Intact | 5 (4.0%) | 121 (96.0%) | 126 |
| Sum | 56 | 129 | 185 |

**Table 5.** Detection results of suspected area of perforation when IoU > 0.5 was defined as valid.

| Ground truth | Prediction (IoU > 0.5) |  | Sum |
| --- | --- | --- | --- |
|  | Suspicious perforation | Intact |  |
| Suspicious perforation | 48 (81.4%) | 11 (13.6%) | 59 |
| Intact | 5 (4.0%) | 121 (96.0%) | 126 |
| Sum | 53 | 132 | 185 |

### Examples of incorrect detection

With the reference set to an IoU score of greater than 0.5, where the tympanic membrane detection was 100% achieved, successful detection and segmentation for both the malleus with its side (right vs left) and the suspected perforation area were achieved in 82.7% (153/185), while neither of the structures were correctly recognized in only 1.6% (3/185). Exemplary images with incorrect detection results are shown in **Fig 5**. The malleus was not properly identified (**Fig 5A**); healed perforation was recognized as a pathologic lesion (**Fig 5B**); and ventilation tube was recognized as a pathologic lesion (**Fig 5C**).

**Fig 5.**
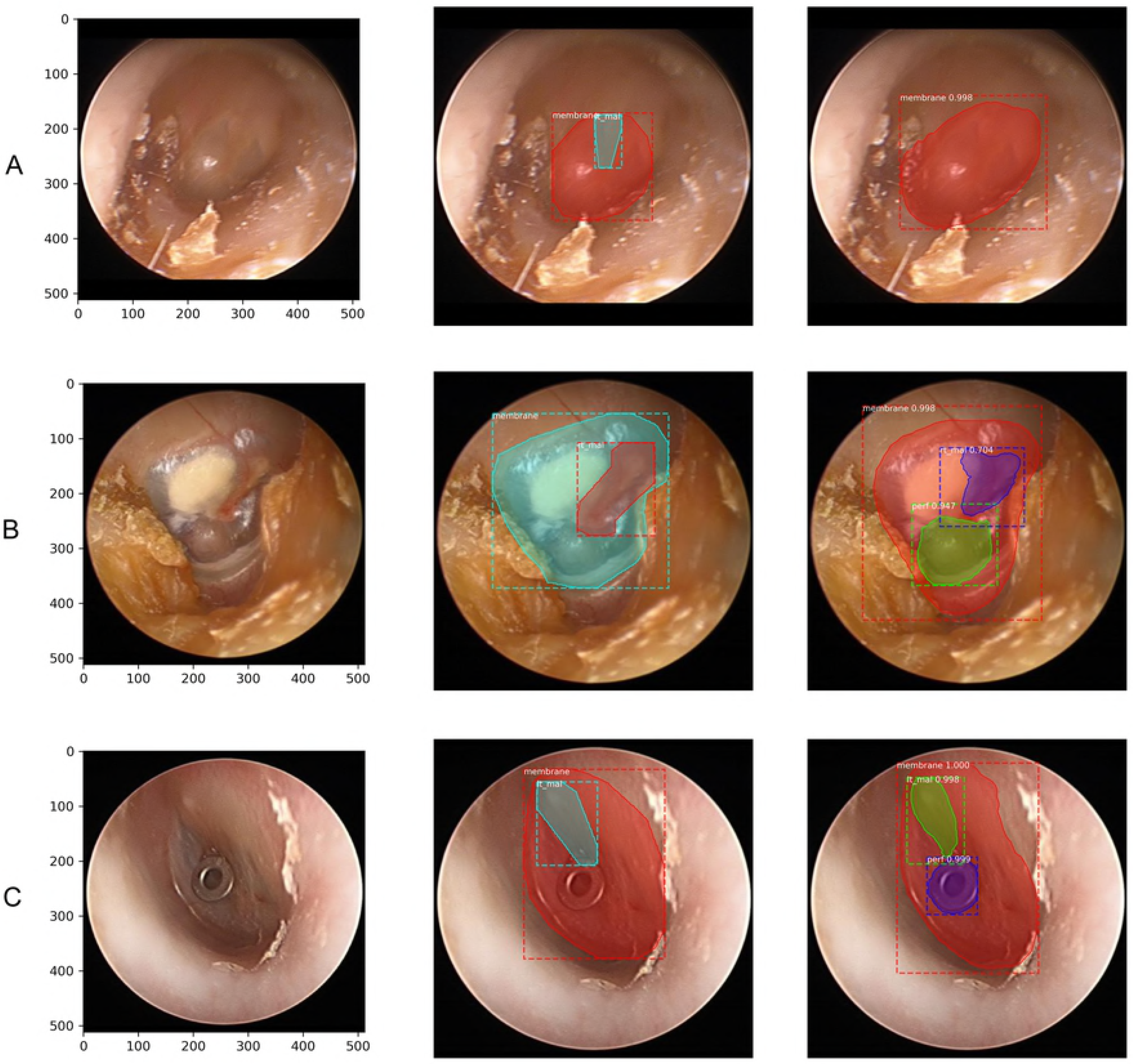
Exemplary images with incorrect detection results. Original images (left), ground truth (center) and outputs (right). The malleus is not properly identified (A), healed perforation is erroneously recognized as pathologic lesion (B), and ventilation tube was recognized as pathologic lesion (C).

### Verification of the versatility of the model through external image

Verification results of our model using google-searched, random images are shown in **S4 Fig**. Although the results were not perfect, the goal of providing a universal model that can be implemented throughout various user environments was achieved.

## Discussions

Visual examination by experienced otoscopists is the most important element in making a diagnosis of AOM or OME [1]. Previously, the diagnostic accuracy of the visual examination by doctors from three countries were measured using a video-recorded otoscopic examination, and the accuracy of the examination by otolaryngologists turned out to be superior to that by pediatricians and general practitioners in all three countries [1].

The relatively poor accuracy may imply that a proper referral to ear specialists cannot be made in many cases, consequently resulting in improper treatment. Thus, ancillary tools may be needed to improve the accuracy, and a good example of such tool has recently been published in the field of ophthalmology. By using the deep learning techniques in three-dimensional optical coherence tomography (OCT) scan data, a novel framework was developed aiding non-specialists to make referral suggestions comparable to clinical experts [10]. Likewise, the development of an auxiliary tool ensuring a certain level of diagnostic accuracy comparable to that by the tympanic endoscopy specialist would provide patients with optimal care.

Until now, most of the deep learning studies using medical images have mainly focused on X-ray, CT, and MRI images, regardless of the target organs [11]. The next tiers of prevalent areas that drew attention of deep learning studies are microscopy and fundus examination [11].

Since ENT practice relies on visual examination based on 2D images, such as tympanic endoscopy and laryngoscopy, there may be greater applicability of the deep learning technology than other fields. However, to the best of our knowledge, there have only been a few studies evaluating deep learning in the field of ENT.

Instead, there have been a couple of studies employing an algorithm-based, automated diagnostic software. Specifically, back in 2013 when the deep learning technology was not widely used, Kuruvilla et al. extracted the features of TM and classified them using the support vector machine (SVM) algorithm; they showed 89.9% accuracy in diagnosing AOM, OME, and no effusion [5]. In 2016, another study showed an accuracy of 80.6% in distinguishing obstructive external auditory canal, normal TM, AOM, OME, and COM using the feature extraction and decision tree algorithm [12]. Recently, a few studies using deep learning have been published; one study revealed 84.4% of accuracy in distinguishing abnormal tympanic membrane from normal tympanic membrane [3], and another study reported 82.2% of accuracy in distinguishing normal from AOM [4]. It is difficult to determine the degree of accuracy and reliability of deep learning-based diagnosis, because its accuracy was not compared directly with that of tympanic endoscopy by a non-ENT doctor; however, from a clinical perspective, a binary classification system – as either abnormal or normal findings – has a significant clinical limitation.

To make breakthroughs, we aimed to establish a model where detection and segmentation of the structure of the tympanic membrane can be performed accurately. To accomplish this and yield objectively interpretable results, we analyzed each anatomical structure with clinical significance rather than analyzing the whole image.

In this study, the Mask R-CNN [6] with ResNet-50 [7] backbone – which was further modified to fit for our data – was used, and image augmentation was implemented to overcome the limited number of training images. Structures with little clinical significance were excluded during the study, while the tympanic membrane, malleus, and its orientation and suspected perforated area were specifically targeted.

As a result, the model was able to detect 100% of the TM, when an IoU score exceeded 0.5, indicating a good prediction in semantic segmentation. Moreover, the model succeeded in the detection of 95.1% of TM even with an increased IoU criterion of 0.7. The result of evaluating the side of the tympanic membrane through the malleus segmentation showed 88.6% accuracy and 99.2% PPV at an IoU score of greater than 0.5. The accuracy of segmentation of suspected perforated area was 91.4% at an IoU score of greater than 0.5.

These high detection rates and accurate segmentations may provide a theoretical basis for effective deep learning training in future studies, by utilizing the segmented area as an input and discarding unnecessary areas. On top of this, it seems very promising that our model is practical and applicable clinically. The visual segmented images offered not only a distinction between normal and abnormal findings, but also an explanation about the pathologic findings. Furthermore, the recognition of these structures will be a starting point for further developing an automated machine providing detailed descriptions of the tympanic membrane findings to be easily included in the electronic medical records.

When an IoU score of greater than 0.5 was used as the reference point, the accuracy of recognizing the TM or perforation area was higher than that of recognizing the malleus. This observation was probably caused by the difference in the size of the object and sharpness of the border. Indeed, in endoscopy images, the malleus seemed small, and the border was not clear compared with other structures. Improved algorithms may be needed to improve the accuracy either by sharpening the border or by improving the accuracy of the method of defining the shape of the malleus.

The results of this study in shown in **Fig 5B**, and our model was able to indicate an equivocal, error-prone lesion, similar to the diagnosis made by experts. Specifically, the healed perforation would not be diagnosed as perforation by experts; however, it can be frequently considered as perforation by an inexperienced individual. The result shown in **Fig 5B** was not what we had initially expected; however, when considering the fact that this deep learning model was developed to play an auxiliary role in the screening tool, our result may be meaningful in terms of increasing sensitivity. In further studies, defining stricter criteria for the detection of pathologic lesions may allow greater accuracy in labeling.

There are several limitations of this study. First is the small set of images included for analysis. Although the image augmentation method was used and obtained meaningful results, we believe that the more images would have yielded greater accuracy. In **Fig 5C**, the result that ventilation tube was recognized as a perforation was caused undoubtedly by the lack of data for ventilation tube. Perforation and healed perforation can also be distinguished via the training of a larger number of images. The second limitation is that the data used are still images, not a video. In real practice, medical decision is often made based on consecutive, serial images rather than still images. In other words, the initially unseen or unclear findings could become visible or clearer during continuing examination. In other words, 82.7% of accuracy obtained in the detection of major three target objects in this study could have been higher if the analysis had been performed based on superimposed continuous images. The third limitation is that parameter tuning was not done. Generally, this process is performed to improve accuracy; however, we concluded that it was not significant because the final goal is to detect and diagnose lesions in a moving image or video, not in still images as described above.

## Conclusion

To the best of our knowledge, this is the first study regarding the semantic segmentation of the tympanic membrane structures and its pathologic findings using deep learning model, demonstrating its clinical applicability. Through the Mask R-CNN and image augmentation, effective machine learning was achieved. Accuracy of structure detection was over 90% when an IoU score of greater than 0.5 was used as the reference. Anatomical segmentation may allow the inclusion of an explanation provided by deep learning as part of the results. This method is applicable not only to tympanic endoscope, but also to sinus endoscope, laryngoscope, and stroboscope. Finally, it will be the starting point for the development of automated medical records descriptor of endoscope images.

## Acknowledgement

This research was supported by Basic Science Research Program through the National Research Foundation of Korea (NRF) funded by the Ministry of Education (2018R1A2B2001054) and a grant of the Korea Health Technology R&D Project through the Korea Health Industry Development Institute (KHIDI), funded by the Ministry of Health & Welfare, Republic of Korea (grant number: HI15C1632, and HI17C0952).

## Supporting information

**S1 Fig. The target structures to be segmented.** (A) The tympanic membrane (TM), malleus with orientation, suspicious perforation, cone of light, cerumen, and TSP, (B) but cone of light, cerumen, and TSP are excluded during the study. Note that 20 pixel dilation of originally drawn mask was used in the final model. (H: height, W: weight, rt_mal: right malleus, col: cone of light)

**S2 Fig. The original image (top left) is augmented by 50 images, using horizontal flip, noise addition, blurring, rotation, and zoom in and out methods.**

**S3 Fig. By eliminating unnecessary areas, memory loss can be prevented and learning efficiency can be improved.** Original image (A) is processed using Otsu binary thresholding (B) and adaptive thresholding (C). By summing the results of the two regions, we can obtain the coordinates of the area we need to obtain (D). The image is obtained by using the coordinates (E), and then the long-axis is transformed to 512 pixels by maintaining the aspect ratio, and then the 512 × 512 image is obtained by using the padding (F).

**S4 Fig. The output results of the model using google-searched images.** Screen captured while searching for images on Google (left) and the result of using that image (right). Note: both tympanic membrane images by Michael Hawke, MD, retrieved from https://upload.wikimedia.org/wikipedia/commons/474c/Traumatic_Perforation_of_the_Tympanic_Membrane.jpg (A) and https://upload.wikimedia.org/wikipedia/commons/1/15/Subtotal_Perforation_of_the_right_tympanic_membrane.tif (B) used under Creative Commons Attribution 4.0 International license.

