## Supplementary figures and images for "The semantic segmentation approach for normal and pathologic tympanic membrane using deep learning"

### Supplemental Figures

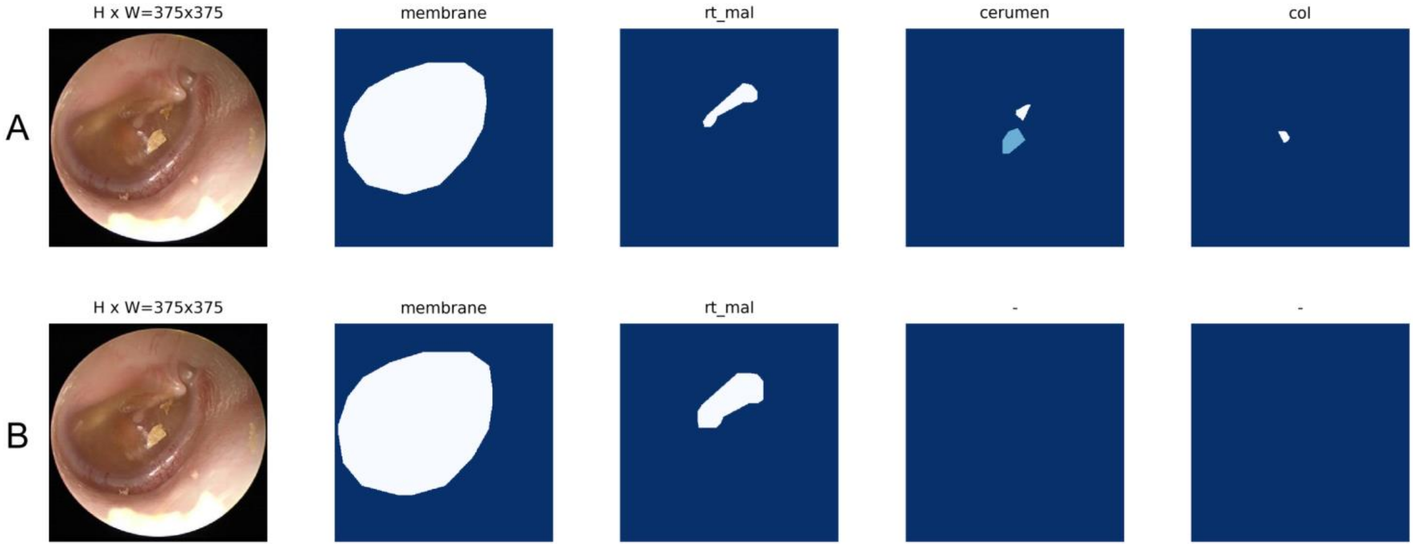

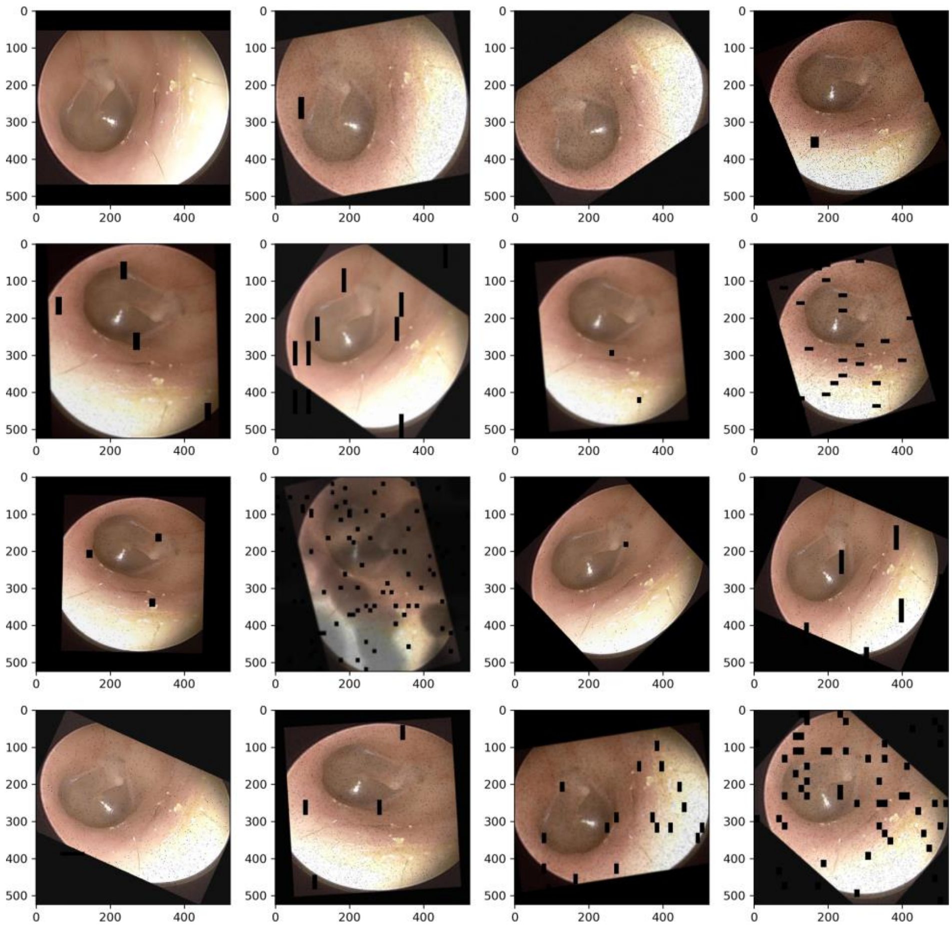


S2 Fig

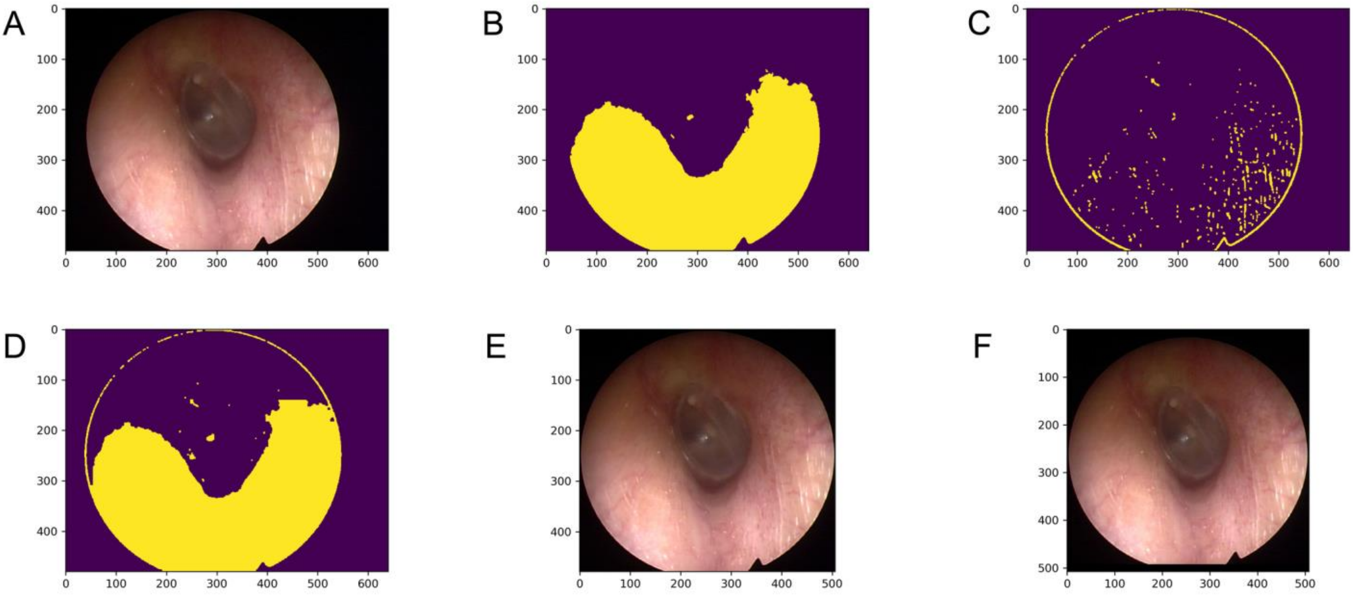


S3 Fig

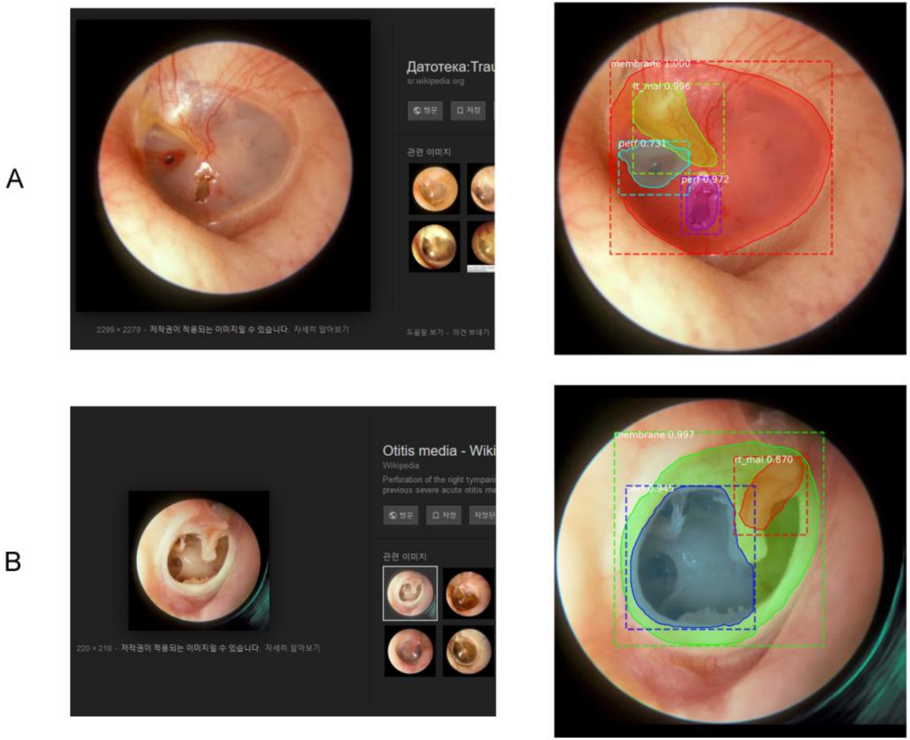
